# Cytoskeleton Regulates Cytoophidium Dynamics in *Drosophila* Ovaries

**DOI:** 10.1101/2025.09.19.677342

**Authors:** Xiao-Jing Liu, Yi-Lan Li, Shu-Yu Pang, Ji-Long Liu, Kun Dou

## Abstract

Cytoophidia are filamentous structures composed of CTP synthase (CTPS) and were initially discovered in ovarian cells of *Drosophila*. As a highly conserved membraneless organelle present across all three domains of life, cytoophidia display dynamic behaviors that are crucial for cellular homeostasis and function. Previous research has shown that cytoophidia are actively transported from nurse cells to oocytes, implicating their potential role in *Drosophila* oogenesis. Nevertheless, the molecular and cellular mechanisms governing cytoophidium dynamics remain largely unclear. In this study, we employ live-cell imaging to systematically characterize the spatiotemporal dynamics of cytoophidia and to explore the underlying regulatory mechanisms. Our findings demonstrate that cytoophidium dynamics are dependent on the cytoskeletal components— microtubules, microfilaments (actin filaments), and myosin II. Notably, disruption of either microtubules or microfilaments resulted in the disassembly or depolymerization of large cytoophidia (macro-cytoophidia), highlighting an essential role of the cytoskeleton in maintaining cytoophidium integrity and assembly. Together, these results indicate that microtubules, microfilaments, and myosin II play pivotal roles in regulating cytoophidium dynamics. This study provides new insights into the mechanisms underlying cytoophidium transport and assembly, and lays a foundation for further investigation of their functional significance in *Drosophila* oogenesis.

## INTRODUCTION

Cytidine-5′-triphosphate (CTP) is a crucial nucleotide that plays essential roles in nucleic acid synthesis, phospholipid metabolism, and energy transfer. The enzyme CTP synthase (CTPS) catalyzes the rate-limiting step in the de novo biosynthesis of CTP [1, 2]. In May 2010, CTPS was unexpectedly found to assemble into elongated, filamentous structures termed cytoophidia in *Drosophila* ovarian cells [3]. Shortly thereafter, similar filamentous CTPS assemblies were independently reported in the bacterium *Caulobacter crescentus* [4] and the yeast *Saccharomyces cerevisiae* [5]. Since these pioneering discoveries, cytoophidia have been identified in a wide range of organisms, including human cells [6], fission yeast [7], plants [8], and archaea [9], underscoring their remarkable evolutionary conservation across the three domains of life.

Although the precise biological functions of cytoophidia remain incompletely understood, emerging evidence suggests their involvement in a variety of cellular processes, such as protein homeostasis and lifespan regulation [10], developmental switch [11], cell adhesion [12], brain development [11], maintenance of cell polarity [13], Intracellular transport [14], immune cell differentiation [15], oncogenesis [16], stress responses [9], and extracellular matrix regulation [17]. These findings point to the potential involvement of cytoophidia in diverse aspects of cellular physiology and adaptation.

In *Drosophila* ovaries, cytoophidia are present in all three major cell types: nurse cells, oocytes, and follicle cells. The polymerization of CTPS into cytoophidia has been shown to support proper egg development, an effect that becomes particularly evident when flies are exposed to the antimetabolite DON (6-diazo-5-oxo-l-norleucine), an inhibitor of CTP synthesis [18, 19]. This observation highlights the important contribution of cytoophidia to oogenesis and suggests a functional link between CTP metabolism and cytoophidium integrity.

Furthermore, our previous work has demonstrated that cytoophidia in the *Drosophila* ovary are capable of active transport from nurse cells to oocytes through intercellular ring canals [20]. This intercellular trafficking is believed to facilitate the transfer of essential biomolecules required for oocyte growth. However, the molecular and cellular mechanisms that govern the directional transport and dynamic behavior of cytoophidia remain poorly defined.

In multicellular organisms, the precise transport and spatial organization of organelles are essential for normal cellular development and the execution of physiological functions [21]. A *Drosophila melanogaster* ovarian egg chamber consists of 16 interconnected germ cells—including 15 nurse cells and 1 oocyte—surrounded by a monolayer of somatic follicle cells [22]. Oogenesis in Drosophila is divided into 14 morphologically distinct stages [23]. Throughout these stages, the oocyte remains transcriptionally quiescent [24] and depends heavily on nurse cells for the supply of critical components such as mRNAs [25], proteins, and organelles [26], which are delivered through specialized cytoplasmic bridges known as ring canals. This dynamic intracellular transport system facilitates the exchange of essential materials and makes the *Drosophila* ovary an ideal model for studying the mechanisms of membraneless organelle trafficking.

Microtubules and actin filaments are two fundamental cytoskeletal components in all eukaryotic cells[27]. They play essential roles in multiple critical cellular functions, including cell division, cell migration, cargo transport and polarity establishment[28–31]. Microtubule-based transport is mediated primarily by two types of motor proteins—kinesins and dyneins—that move cargoes toward the plus-end and minus-end of microtubules, respectively, according to their intrinsic polarity [32–34]. Kinesin family motors generally transport cargoes toward the microtubule plus-end, while cytoplasmic dynein moves toward the minus-end. Actin-based motility, on the other hand, is driven predominantly by myosin motor proteins that move along actin filaments [35].

While the roles of the cytoskeleton in general intracellular transport are well established, its specific involvement in the regulation of cytoophidium motility and assembly remains largely unexplored. Understanding how cytoskeletal elements interact with cytoophidia may provide critical insights into the mechanisms underlying their transport and functional integration within cells.

In this study, we utilize live-cell imaging to investigate the dynamic behavior of cytoophidia in *Drosophila* ovaries under physiological conditions. Our real-time imaging data reveal that the cytoskeleton—particularly microtubules—plays a pivotal role in regulating cytoophidium dynamics. In control samples, cytoophidia exhibit robust motility, including active directional transport from nurse cells to oocytes. However, upon disruption of microtubules, we observe a significant reduction in cytoophidium movement and a loss of oocyte-directed trafficking, suggesting an essential requirement for the microtubule network in cytoophidium assembly and transport.

## MATERIALS AND METHODS

### Fly Stocks

All stocks were maintained at 25 °C on standard cornmeal. The following fly stocks were used in this study: Both *w^1118^* and C-terminal mChe-4V5 tagged CTPS knock-in flies out of *w^1118^* produced in our laboratory were used as wild-type controls unless stated otherwise. The stocks used were: *Sp/Cyo; Sb/Tm6B* (Institute of Biochemistry and Cell Biology, Chinese Academy of Sciences, *Drosophila* Resources and Technology Platform), all stocks were maintained at 25 °C on standard cornmeal.

The GAL4/UAS system (Brand and Perrimon, 1993) was utilized for adipocyte-specific expression or RNAi knockdown of the desired genes. All RNAi stocks were from the TRiP collection (Bloomington). The stocks used were: *UAS-khc-RNAi* (Bloomington *Drosophila* Stock Center # 41858); *UAS-Rab11-RNAi* (Bloomington *Drosophila* Stock Center # 27730); *UAS-Rab6-RNAi* (Bloomington *Drosophila* Stock Center # 27490), *Sp/Cyo; GFP-Pav /Tm6c.sb* (Bloomington *Drosophila* Stock Center # 81657)

The following fly stocks were generated in this study: *nosgal4/Cyo; CTPS^mCherry-KI^/Tm6c.sb*

### Quantitative RT-PCR

Total RNAs were prepared from the lysate of larval fat body tissues. cDNAs were synthesized with PrimeScript RT Master mix (ABclonal) followed by adding template RNA. 2X SYBR Green PCR Master Mix was purchased from ABclonal. Real-time quantitative PCR was conducted using the QuantStudion^™^ 7 flex System (Applied Biosytstems). For normalization, *rp49* was utilized as the internal control. The oligonucleotide primers used were as follows:

*rp49*: sense 5’-TACAGGCCCAAGATCGTGAA-3’, antisense 5’-TCTCCTTGCGCTTCTTGGA-3’.

*CTPS*: sense 5’-GAGTGATTGCCTCCTCGTTC-3’, antisense 5’-TCCAAAAACCGTTCATAGTT-3’.

*Khc:* sense 5’-TGGAGGATCTCATGGAGGCA-3’, antisense 5’-ATGCGCTTCTTCTGGTGTGA-3’.

*Gal4*: sense 5’-CAACTGGGA*GTGTCGCTACTC*-3’, antisense 5’-CACCGTACTCGTCAATTCCAAG-3’.

### Inhibitor treatment

Briefly, 2 - 4day female files raised at 25°C on a standard corn-meal agar diet were starved for 12 h, then these flies were fed with yeast paste supplemented with 200μg/ml colchicine (Adamas, 013456239) for 24h to disrupt tubulin and paclitaxel 250µM for 24h to Stabilize microtubule structure, flies were fed with yeast paste supplemented with Cytochalasin D 500 µM for 24h to disrupt microfilament and para-nitroblebbistatin 500 µM for 24h to inhibit myosin. To inhibit dynein flies were fed with yeast paste supplemented with 200µM Ciliobrevin D for 24h.

### Immunohistochemistry

Egg chambers from stage 7-8 were imaged and quantified. Ovary from flies were dissected in Grace’s Insect Medium (Sigma-Aldrich) and then fixed in 4% formaldehyde (Sigma) diluted in PBS for 10 min before immunofluorescence staining. The samples were then washed twice using PST (0.5% horse serum + 0.3% Triton × 100 in PBS). Samples were incubated with primary antibodies at room temperature overnight and then washed using PBST. Secondary antibodies were used to incubate the samples at room temperature for another night.

Primary antibodies used in this study were rabbit anti-CTPS (1:1000; y-88, sc-134457, Santa Cruz Bio Tech Ltd., Santa Cruz, CA, USA), Secondary antibodies used in this study were anti-mouse, rabbit, or with Cy5 (Jackson ImmunoResearch Laboratories, West Grove, PA, USA). Hoechst 33342 was used to label DNA, Membrane were stained with Rhodamine-conjugated phalloidin (1:5000; SB-YP0059-50T, share-bio). Ovaries were dissected in 1X PBS and fixed 20 min on the rotator in 1X Brinkley Renaturing Buffer 80 (BRB80), pH 6.8 (80 mM piperazine-N, N’-bis (2-ethanesulfonic acid) [PIPES]), 1 mM MgCl2, 1 mM EGTA) +0.1%Triton X-100 +8% EM-grade formaldehyde. After fixation, the samples were briefly washed five times with 1X PBTB and subsequently incubated overnight at 4 °C with FITC-conjugated β-tubulin antibody (ProteinTech, Cat# CL488-66240) at a dilution of 1:100. The ovary was stained with CellMask Green Actin Tracking Stain (Cat. No. A57243) at 1X concentration for 30 mins at 37℃.

### Live imaging of Drosophila egg chamber

A group of 5–15 female flies 2–4 days after eclosion, together with several male flies, were transferred into a fly bottle and a wet yeast paste and incubated at 25 °C for about 1–2 days, and then dissected in 3 mL Grace’s Insect Medium, producing a large number of Stages 6–8 egg chambers.The dissected ovaries were transferred into a 35 mm glass bottom dishes with a 200 uL pipette. Fluorescent samples were imaged using Leica sp8 confocal microscope with 40×oil lens.

### Microscopy and Image Analysis

Fluorescent images were obtained using confocal laser scanning microscopy (Leica SP8) at 20 X, 40 X, or 63 X for oil objects. Images were acquired every 1 μm/step for whole ovariole imaging or 0.3∼0.5μm/step for individual egg chambers in z stacks. We counted the length and number of CTPS cytoophidia in cells by analyzing 40 X confocal images using FIJI-ImageJ. The data were normalized by the number of cells in one image. The resulting binary images of the outlines were then dilated to 8 pixels to create membrane masks in FIJI-ImageJ. All images were obtained under laser-scanning confocal microscopy (Leica SP8 STED 3X) and Nikon TI2-E+CSU W1 Sora 1Camera. ImageJ was used to analyze the area and number of follicle cells. We used ImageJ cell counter to calculate different shapes of cells to get the number of polygonal cells. We used ImageJ to measure the area of each cell. For each statistical quantification.

### Western Blotting

Female adult ovaries of *Drosophila* were collected with gathered into lysis buffer RIPA (Meilunbio, Dalian, China) with protease inhibitor cocktail (Bimake, Shanghai, China) for Western blotting, and then ground with 1 mm Zirconia beads in Sonicator (Shanghai Jing Xin, Shanghai, China). The sample would then lysis on ice for up to 30 min. Samples were centrifuged for 10 min at 10,000 g at 4 °C. The 6× protein loading buffer was pipetted into the supernatants and boiled at 99 °C for 15 min to obtain protein. Then, the protein sample was run through 10% SDS-PAGE gels and transferred to PVDF membranes. At room temperature, membrane was incubated with 5% *w*/*v* nonfat dry milk dissolved by 1× TBST for 1 h of blocking. Then, the membrane was incubated with primary antibodies in 5% *w*/*v* nonfat milk at 4 °C and gently shaken overnight.

The following primary antibodies were used in this study: anti-mCherry Tag Monoclonal antibodies (Cat. No. A02080, Abbkine, Beijing China), mouse anti-a-Tubulin antibodies (Cat. No. T6199, Sigma). The membranes were washed three times for 5 min per time with shaking, then incubated with secondary antibodies (anti-mouse IgG, HRP-linked antibody, Cell Signaling, Danvers, MA, USA) diluted in 5% milk at room temperature for 1 h. An Amersham Imager 600 (General Electric, Boston, MA, USA) and Pierce ECL Reagent Kit (Cat. No. 32106, Thermo Fisher, Waltham, MA, USA) were adopted for the chemiluminescence immunoassay. Protein levels were quantified on ImageJ (National Institutes of Health, Bethesda, MD, USA) and normalized to tubulin. At least three biological replicates were quantified.

### Data Analysis

All data are presented as the mean ± standard error of the mean (s.e.m) from at least three independent experiments. Statistical analysis was performed using unpaired two-tailed Student’s t-test with GraphPad Prism8.0. P < 0.05 was considered to be statistically significant.Images collected by confocal microscopy were processed using Adobe Illustrator and ImageJ. Quantitative analysis was processed by Excel and GraphPad.

## RESULTS

Cytoophidia exhibit three distinct modes of dynamic behavior To investigate the dynamic properties of cytoophidia, we performed live-cell imaging in *Drosophila* ovarian cells using a CTPS-mCherry fusion construct. Our analysis revealed that cytoophidia display three primary types of dynamic behavior within nurse cells and germline cells (including both oocytes and follicle cells): intracellular transport, intercellular movement, and morphological remodeling (**Figure 1**; **Video S1**).

**Figure 1.**
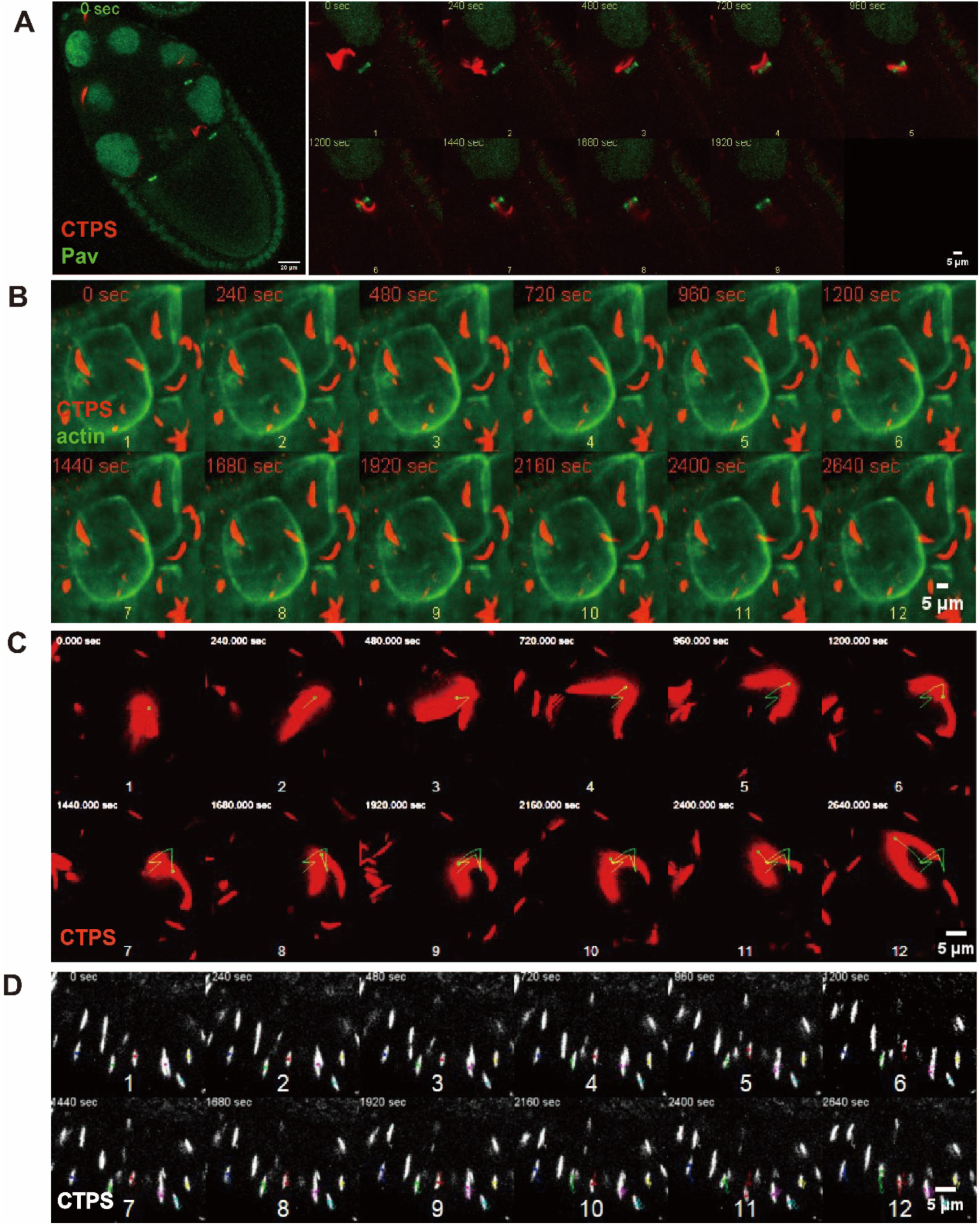
Three types of dynamic changes in cytoophidia. A. Live-cell imaging of cytoophidium transporting from nurse cell to oocyte. CTPS is tagged with mCherry, ring canal labeled by GFP-Pav. Time stamps are on the top-left, images are taken for every 4 minutes. Scale bar, 20μm (left), 5μm (right). B. Live-cell imaging of cytoophidium transporting from one nurse cell to the other nurse cell. CTPS is tagged with mCherry, actin labeled by CellMask Green Actin Tracking Stain. Time stamps are on the top-left, images are taken for every 4 minutes. Scale bar, 5μm. C. Trajectories of cytoophidia in germline cells. CTPS is tagged with mCherry. Time stamps are on the top-left, images are taken for every 4 minutes. Time stamps are on the top-left, images are taken for every 4 minutes. Scale bar, 5μm. D. D. Trajectories of cytoophidia in follicle cells. CTPS is tagged with mCherry. Time stamps are on the top-left, images are taken for every 4 minutes. Time stamps are on the top-left, images are taken for every 4 minutes. Scale bar, 5μm.

Within follicle cells, cytoophidia typically maintain a rigid and linear morphology. Trajectory analyses demonstrated that these cytoophidial structures exhibit bidirectional, back-and-forth motility along the apical-basal axis, suggesting a relatively constrained and oscillatory movement pattern (**Figure 1D**). In contrast, cytoophidia in germline cells (oocytes and nurse cells) show more dynamic and less directionally constrained intracellular movements (**Figure 1A, C**). This increased mobility may facilitate interactions among cytoophidia, potentially contributing to their assembly or coalescence.

Moreover, cytoophidia in germline cells frequently undergo active morphological remodeling, including processes such as twisting, flipping, and bending, often occurring concurrently with their movement. These structural changes further highlight the dynamic nature of cytoophidia in the germline and suggest a high degree of structural plasticity during their intracellular behavior (**Figure 1A, C**).

To specifically visualize the intercellular transport of cytoophidia between nurse cells and oocytes, we employed GFP-Pav to label the ring canals—specialized cytoplasmic bridges that connect these cells (**Figure 1A**). Time-lapse imaging clearly captured instances of direct cytoophidium transport through ring canals, not only from nurse cells to oocytes but also between adjacent nurse cells (**Figure 1A, B**). Given that the oocyte is transcriptionally silent during much of oogenesis and relies heavily on nurse cells for the supply of essential biomolecules, including organelles, proteins, and mRNAs, this intercellular trafficking likely plays a critical role in oocyte development. Accordingly, we hypothesize that the directed movement of cytoophidia through ring canals may contribute to the delivery of functional components that support oocyte maturation.

### Microtubules disassembly impairs cytoophidium dynamics and assembly

In eukaryotic cells, intracellular transport is primarily mediated by motor proteins that move along two major cytoskeletal networks: microtubules and actin filaments. To identify the cytoskeletal component(s) responsible for driving the dynamic behavior of cytoophidia, we disrupted each cytoskeletal system individually and examined the resulting effects on cytoophidium dynamics.

We first disrupted microtubule networks by treating adult flies with 200 μg/mL colchicine for 18 hours. The efficacy of microtubule depolymerization was assessed by immunofluorescence staining against α-tubulin or β-tubulin. Compared to the control group, colchicine treatment resulted in a marked disruption of the microtubule cytoskeleton, with barely detectable intact microtubule signals in treated samples (**Figure 2B**).

**Figure 2.**
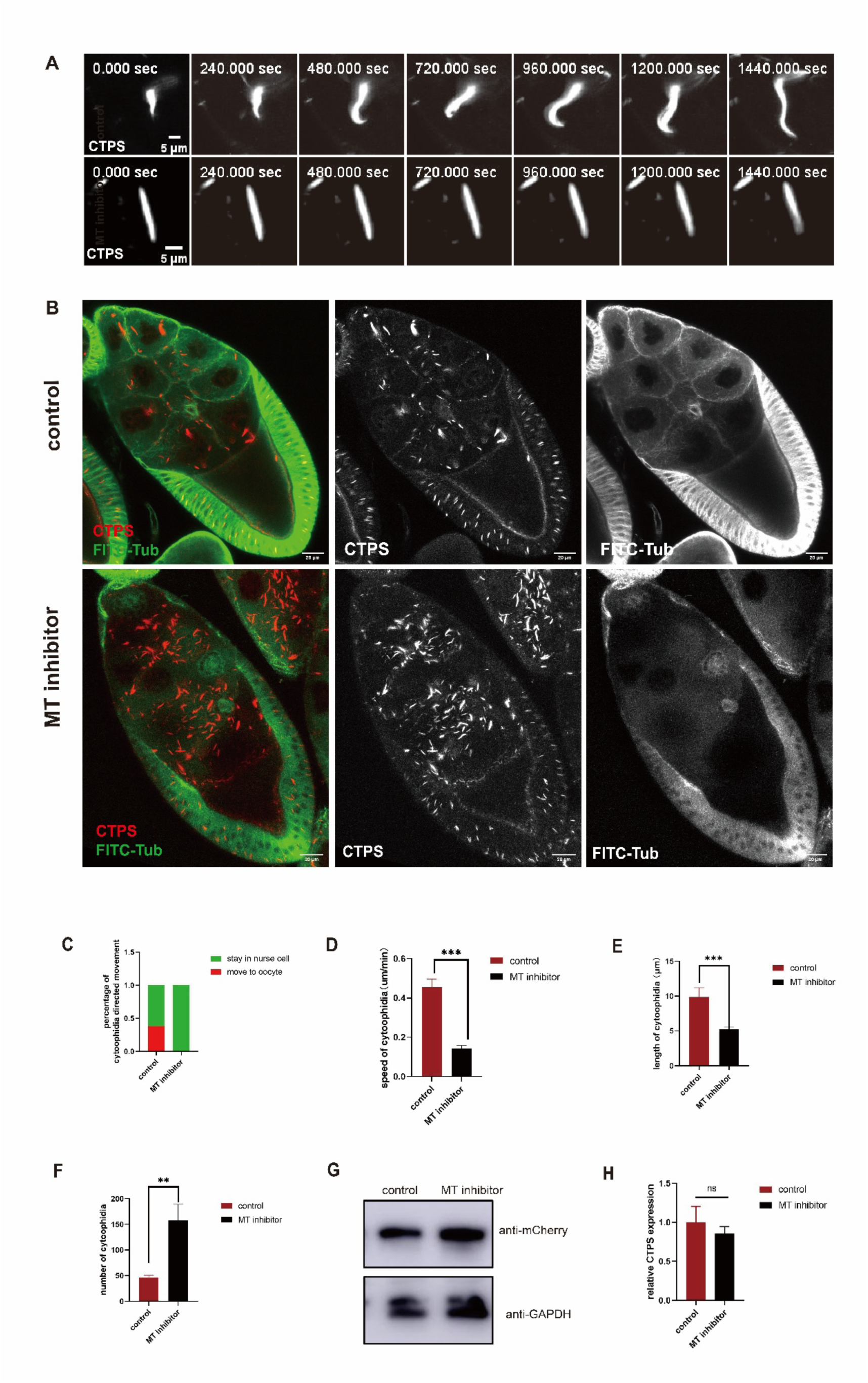
Microtubules are essential for cytoophidia morphological changes movement, and assembly. (A) Time-lapse imaging revealed cytoophidia morphological changes, The time marked in the upper left corner represents the time since the capture began. The CTPS (gray) signal shown is obtained using mCherry-tagged CTPS. Scale bar = 5μm. (B) Cytoophidia abundance and length are affected by microtubule depolymerization. Microtubules (green) are labeled with FITC-conjugated tubulin antibody, CTPS (red) signal is obtained using mCherry-tagged CTPS. scale bars, 20μm. (C) Quantification of cytoophidium intercellular transportation through nurse cell-oocyte ring canals in control and microtubule-inhibited groups. (D) The speed of cytoophidia movement. We analyzed 35 filaments in control group and 35 filaments in microtubule depolymerization group, Data are represented as mean ± SEM. ***, p-value = 0.0001. (E) Length of cytoophidia in ovary nurse cell. We analyzed 32 filaments in control group and 88 filaments in microtubule depolymerization, Data are represented as mean ± SEM. ***, p-value = 0.0001. (F) Number of cytoophidia in ovary nurse cell. We analyzed 9 ovaries in control group and 413 filaments in total, and 9 ovaries in microtubule depolymerization and 1417 filaments in total, Data are represented as mean ± SEM. **, p-value = 0.003. (G) Western blotting analysis of CTPS proteins from ovary of CTPS-mCherry fly. Anti-mCherry antibody was used for the immunoblotting analysis. GAPDH was used as an internal control. (H) Relative CTPS mRNA expression levels are measured by quantitative RT-PCR using ovary from the control and MT inhibitor-treated flies (6 ovaries/group, 3 biological replicates). Data are represented as mean ± SEM. ns, no significance in difference.

Concomitant with microtubule loss, we observed a striking stabilization of cytoophidium morphology: instead of exhibiting the dynamic shape-shifting behavior seen in control cells, cytoophidia in colchicine-treated samples displayed a static, unchanging structure (**Figure 2A; Video S2**). This indicates that intact microtubules are required to sustain the dynamic morphological transitions of cytoophidia.

To further characterize the impact of microtubule disruption on cytoophidium motility, we analyzed their movement trajectories. Strikingly, microtubule depolymerization led to a severe suppression of cytoophidium dynamics, with significantly reduced or absent motility (**Figure 2D; Video S2**). More importantly, directed transport of cytoophidia from nurse cells to the oocyte via ring canals was completely abolished (**Figure 2C**). This finding strongly suggests that functional microtubule networks are essential for the directional, intercellular movement of cytoophidia.

In addition to impaired motility, microtubule disruption also had a notable impact on cytoophidium assembly. Quantitative analysis revealed a significant increase in the number of cytoophidia, accompanied by alterations in their filament lengths (**Figures 2B, 2E, 2F**). To rule out the possibility that this increase was due to elevated CTPS expression, we performed quantitative PCR (qPCR) and Western blotting to measure CTPS transcript and protein levels, respectively. Importantly, neither CTPS mRNA nor protein abundance was significantly altered following colchicine treatment (**Figure 2G, 2H**). These results indicate that microtubules regulate cytoophidium assembly independently of changes in CTPS expression levels.

Taken together, these findings demonstrate that microtubules play a central role in coordinating multiple core aspects of cytoophidium biology in the *Drosophila* germline, including morphological dynamics, intracellular and intercellular motility, and the assembly process.

### Stabilization of microtubules does not significantly alter cytoophidium dynamics or assembly

Having established that microtubule disassembly disrupts cytoophidium dynamics and impairs their assembly, we next sought to investigate whether microtubule stabilization similarly influences these processes. To this end, we treated adult *Drosophila* with the microtubule-stabilizing agent paclitaxel at a concentration of 200 μM for 24 hours.

Immunofluorescence analysis revealed a significant increase in both microtubule fluorescence intensity and polymer density in paclitaxel-treated samples compared to untreated controls, indicative of robust microtubule stabilization (**Figure 3B**). This pharmacological effect confirms the effectiveness of paclitaxel in inducing stable microtubule networks in our experimental system.

**Figure 3.**
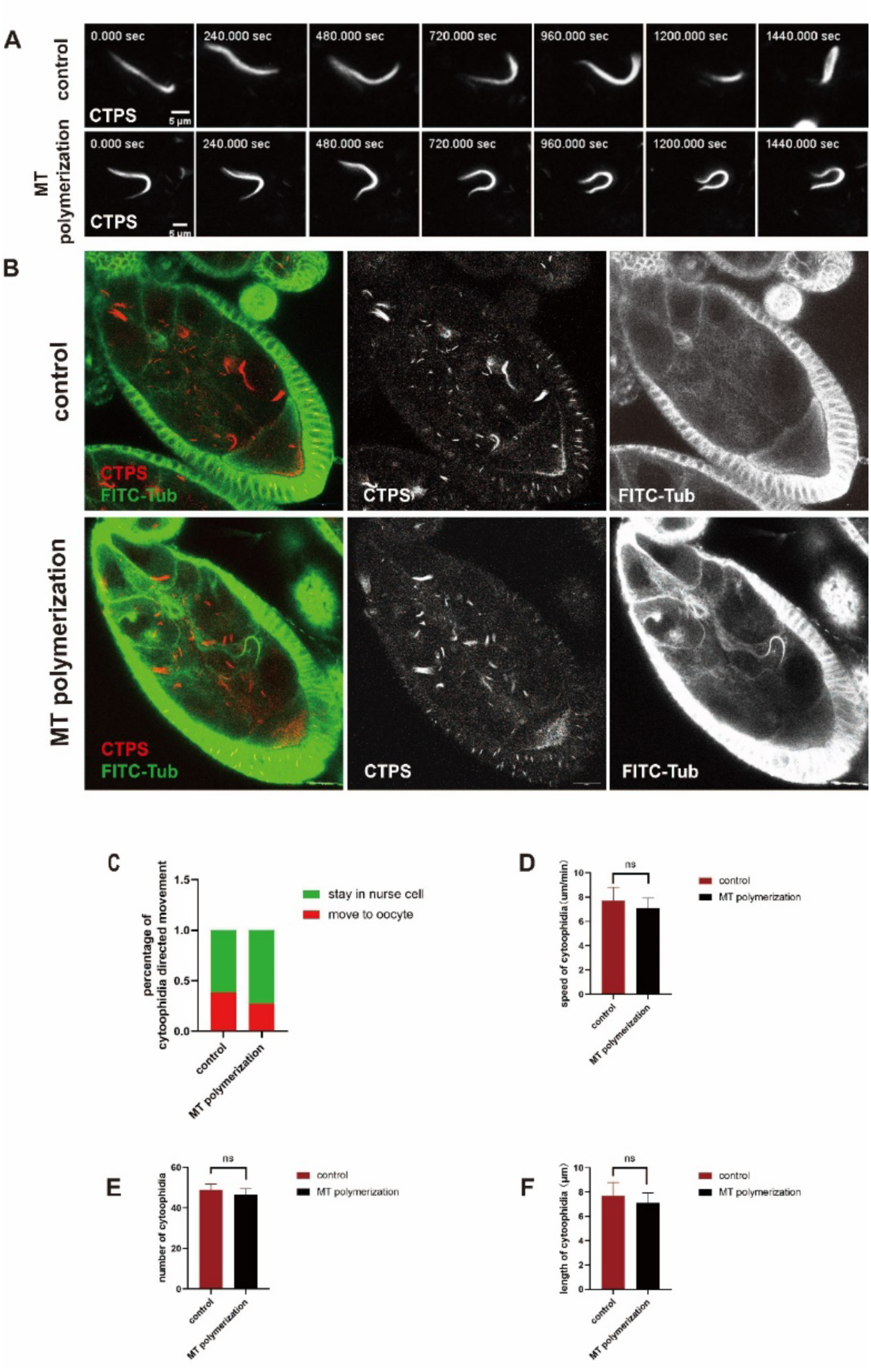
Microtubules stablization does not affect motion, morphological changes and assembly. (A) Time-lapse imaging revealed cytoophidia morphological changes, The time marked in the upper left corner represents the time since the capture began. The CTPS (gray) signal shown is obtained using mCherry-tagged CTPS. Scale bar = 5μm. (B) Cytoophidia abundance and length are not affected by microtubule polymerization. Microtubules (green) are labeled with FITC-conjugated tubulin antibody, CTPS (red) signal is obtained using mCherry-tagged CTPS. scale bars, 20μm. (C) Quantification of cytoophidia transport through nurse cell-oocyte ring canals in control and microtubules polymerization groups. (D) The speed of cytoophidia movement. We analyzed 47 filaments in control group and 47 filaments in microtubule polymerization group, Data are represented as mean ± SEM. ns, no significance in difference. (E) Number of cytoophidia in ovary nurse cell. We analyzed 5 ovaries in control group and 245 filaments in total, and 5 ovaries in microtubule polymerization and 233 filaments in total, Data are represented as mean ± SEM. ns, no significance in difference. (F) Length of cytoophidia in ovary nurse cell. We analyzed 47 filaments in control group and 47 filaments in microtubule polymerization, Data are represented as mean ± SEM. ns, no significance in difference.

To assess the impact of microtubule stabilization on cytoophidium behavior, we performed live-cell imaging under physiological conditions. Strikingly, cytoophidia in paclitaxel-treated flies retained their characteristic morphological plasticity, exhibiting a range of dynamic behaviors similar to those observed in control samples (**Figure 3A; Video S3**). Individual cytoophidial structures continued to display heterogeneous motility patterns, and key intrinsic motility parameters—including speed and directional persistence—remained largely unchanged (**Figure 3D; Video S3**).

Quantitative analysis further demonstrated that neither the number nor the average length of cytoophidia was significantly altered in the paclitaxel-treated group compared to the control (**Figures 3E, 3F**). These findings suggest that microtubule stabilization does not overtly enhance or inhibit the dynamic assembly or maintenance of cytoophidia.

Taken together, our results indicate that while microtubule disassembly disrupts cytoophidium dynamics and assembly, microtubule stabilization alone is insufficient to modulate these processes. We therefore speculate that cytoophidium assembly and dynamics may depend more critically on the polymerization state or net balance of microtubule dynamics (e.g., dynamic instability or “threadmilling”) rather than merely on the presence of stabilized microtubule polymers.

### Cytoplasmic dynein drives the directed trafficking of cytoophidia

Given that microtubules are essential for the directed transport and dynamics of cytoophidia, we next sought to identify the specific motor proteins responsible for mediating this process. Among the known microtubule-based motor proteins, cytoplasmic dynein is a minus-end-directed motor that plays a central role in the intracellular transport of various membranous and membraneless cargoes, including peroxisomes, autophagosomes, lipid droplets, and mitochondria[36]. Given that cytoophidia are membraneless organelles that undergo directed transport from nurse cells to oocytes through microtubule-rich ring canals, we hypothesized that dynein may also be involved in their long-range, directional motility.

To test this hypothesis, we inhibited dynein activity using 500 μM ciliobrevin D, a small-molecule inhibitor that specifically blocks cytoplasmic dynein function [37], and examined its effects on cytoophidium behavior.

Live imaging analysis revealed that dynein inhibition did not significantly affect cytoophidium morphology (**Figure 4A**). Moreover, both the dynamic behaviors and movement trajectories of cytoophidia remained largely unchanged in the presence of ciliobrevin D (**Figures 4B, 4C; Video S4**). These observations suggest that dynein is not essential for maintaining the general motility or morphological plasticity of cytoophidia under these conditions.

**Figure 4.**
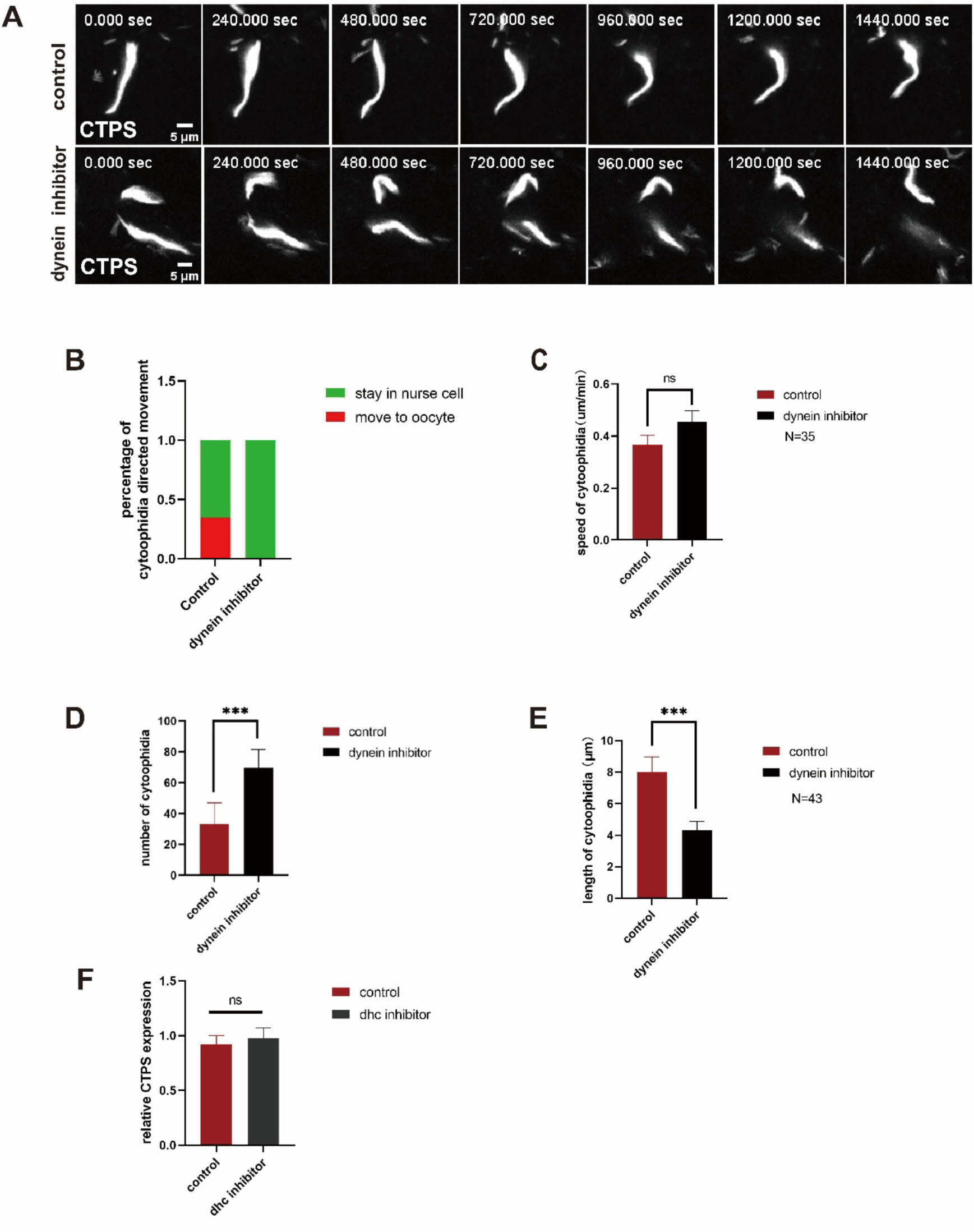
Dynein is indispensable assembly and directed movement of cytoophidia. (A) Time-lapse imaging revealed cytoophidia morphological changes, The time marked in the upper left corner represents the time since the capture began. The CTPS (gray) signal shown is obtained using mCherry-tagged CTPS. Scale bar = 5μm. (B) Quantification of cytoophidia transport through nurse cell-oocyte ring canals in control and dynein inhibited groups. (C) The speed of cytoophidia movement. We analyzed 35 filaments in control group and 35 filaments in dynein inhibited group, Data are represented as mean ± SEM. ns, no significance in difference. (D) Length of cytoophidia in ovary nurse cell. We analyzed 43 filaments in control group and 47 filaments in dynein inhibited, Data are represented as mean ± SEM. ***, p-value = 0.0009. (E) Number of cytoophidia in ovary nurse cell. We analyzed 8 ovaries in control group and 265 filaments in total, and 8 ovaries in dynein inhibited and 557 filaments in total, Data are represented as mean ± SD. ****, p-value = 0.0001. (F) Relative CTPS mRNA expression levels are measured by quantitative RT-PCR using ovary from the control and dynein inhibitor-treated flies (6 ovaries/group, 3 biological replicates). Data are represented as mean ± SEM. ns, no significance in difference.

However, quantitative analysis of cytoophidium populations revealed significant changes in their size distribution. Specifically, we observed a marked increase in the proportion of shorter cytoophidium fragments in dynein-inhibited samples compared to controls (**Figures 4D, 4E**). Notably, this alteration in cytoophidium length distribution was not accompanied by significant changes in CTPS mRNA levels (**Figure 4F**), suggesting that the effect is unlikely to result from altered transcription or translation of CTPS. Instead, these data point to a potential role for dynein in regulating cytoophidium assembly or fragment stability.

Taken together, these findings indicate that while dynein inhibition does not impair the overall morphology, motility, or dynamic behavior of cytoophidia, it does lead to measurable changes in cytoophidium size distribution. This suggests that cytoplasmic dynein is primarily involved in the assembly or maintenance of cytoophidium integrity, rather than in their long-range intracellular transport or dynamic remodeling.

### Kinesin plays a limited role in the regulation of cytoophidium dynamics

Kinesin is a well-characterized plus-end-directed microtubule motor protein that functions as a tetrameric complex, consisting of two heavy chains and two light chains[38]. Through its ATP-dependent conformational changes, kinesin generates mechanical force to transport various cellular cargoes toward the plus ends of microtubules[39]. Given its established role in intracellular transport, we hypothesized that kinesin might also contribute to the regulation of cytoophidium motility or transport.

To investigate this possibility, we examined cytoophidium behavior in genetic backgrounds where kinesin heavy chain (KHC) was knocked down via RNA interference (RNAi). Although KHC knockdown (khc RNAi) led to impaired oocyte development, consistent with the known role of kinesin in oogenesis, live-cell imaging revealed that cytoophidium morphological dynamics remained largely unaffected (**Figure 5; Video S5**). Specifically, time-lapse imaging consistently captured ongoing structural transformations of cytoophidia, indicating that their intrinsic plasticity and remodeling processes were preserved in the absence of KHC.

**Figure 5.**
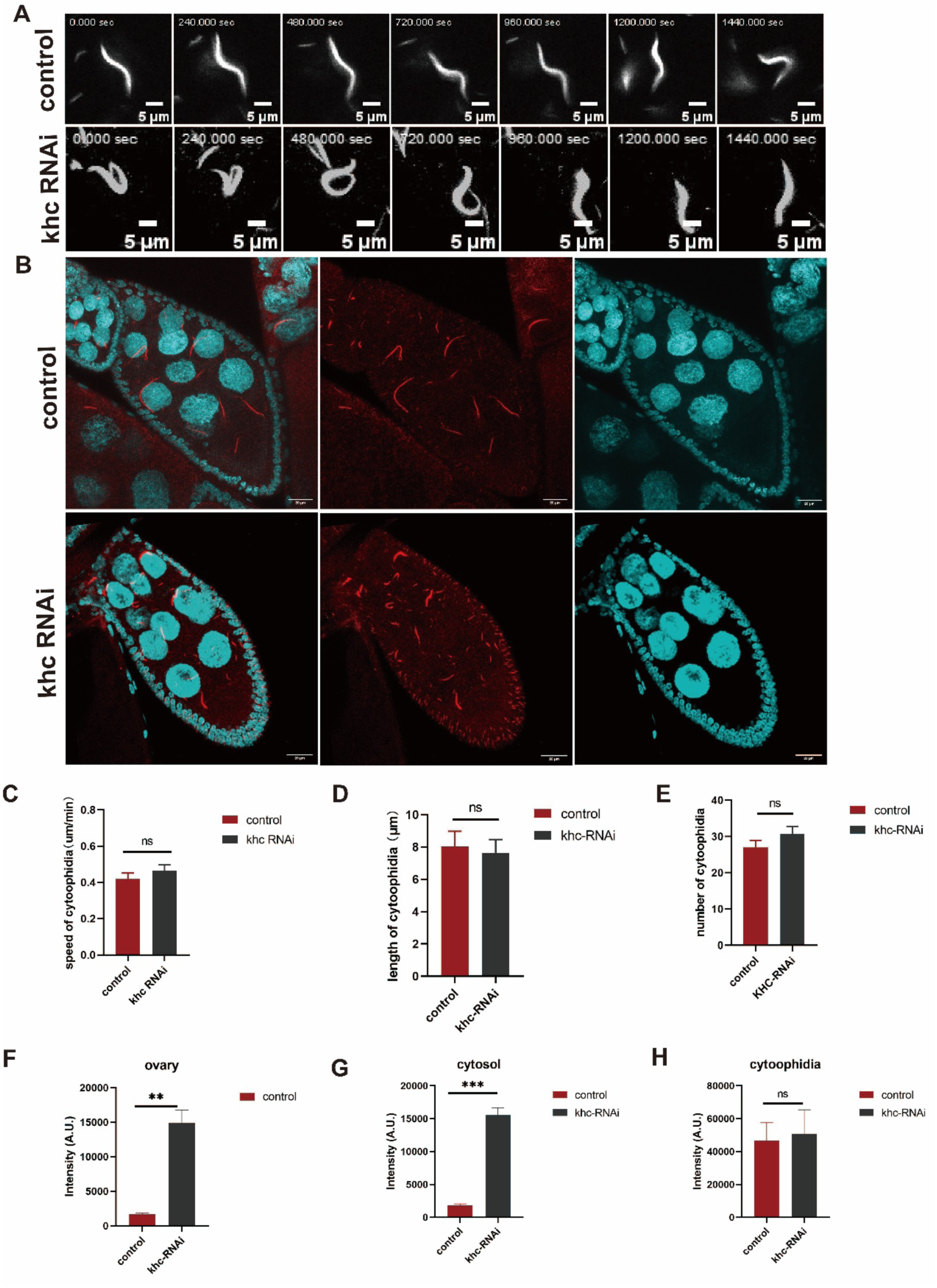
Kinesin does not affect cytoophidia dynamic changes. (A) Time-lapse imaging revealed cytoophidia morphological changes, The time marked in the upper left corner represents the time since the capture began. The CTPS (gray) signal shown is obtained using mCherry-tagged CTPS. Scale bar = 5μm (B) Khc knockdown Drosophila ovaries. CTPS (red)signal shown is obtained using mCherry-tagged CTPS. scale bars, 20μm. (C) The speed of cytoophidia movement. We analyzed 41 filaments in control group and 30 filaments in khc knockdown group, Data are represented as mean ± SEM. ns, no significance in difference. (D) Length of cytoophidia in ovary nurse cell. We analyzed 43 filaments in control group and 44 filaments in khc knockdown, Data are represented as mean ± SEM. ns, no significance in difference. (E) Number of cytoophidia in ovary nurse cell. We analyzed 6 ovaries in control group and 162 filaments in total, and 6 ovaries in khc knockdown and 184 filaments in total, Data are represented as mean ± SEM. ns, no significance. (F-H) Intensity of CTPS-mCherry signal in ovary (**, p-value = 0.0019), cytosol (***, p-value = 0.0002) and cytoophidia (ns, no significance in difference). We analyzed 3 ovaries in control group and 3 ovaries in khc group. Data are represented as mean ± SEM.

Further quantitative analysis of cytoophidium behavior showed that kinesin inhibition did not disrupt their directional transport from nurse cells to oocytes via ring canals. Trajectory analysis confirmed the continued occurrence of long-range, intercellular trafficking, supporting the notion that cytoophidium transport can occur independently of kinesin function.

In addition, no statistically significant changes were observed in the motility rates of cytoophidia, nor was there a noticeable increase in the proportion of fragmented or disassembled cytoophidial structures within the egg chambers of khc RNAi flies.

Interestingly, we observed a significant increase in the overall intensity of CTPS fluorescence in the ovaries of khc RNAi flies compared to control animals. To determine whether this increase was attributable to diffuse, non-filamentous CTPS or to filamentous cytoophidia, we performed a quantitative comparison of mean fluorescence intensity between the two forms in both control and khc RNAi samples. The results revealed that while the intensity of diffuse CTPS was markedly elevated in the khc RNAi background, the fluorescence intensity of cytoophidial structures themselves remained unchanged.

This differential effect suggests that kinesin may be specifically involved in the early assembly or condensation of CTPS into filamentous cytoophidium structures, rather than in the maintenance or dynamics of mature cytoophidia. However, kinesin does not appear to play a major role in the ongoing regulation of cytoophidium motility, transport, or structural stability under physiological conditions.

In summary, these findings indicate that kinesin activity is largely dispensable for the dynamic behavior, intracellular transport, and assembly of cytoophidia in the *Drosophila* ovary. This may reflect functional redundancy among microtubule motor proteins or the existence of kinesin-independent mechanisms for cytoophidium trafficking and assembly.

### The dynamics and assembly of cytoophidia depend on microfilaments

Actin filaments, as key components of the cytoskeletal network, play essential roles in a wide range of cellular processes, including intracellular transport, cell motility, and the maintenance of cellular architecture. Previous studies in the fission yeast *Schizosaccharomyces pombe* have suggested that actin filaments regulate cytoophidium dynamics through the modulation of filament aggregation [14], highlighting a potential conserved role in cytoophidium regulation. To investigate whether this function is evolutionarily conserved in multicellular organisms, we examined the effects of actin filament disruption on cytoophidium behavior in *Drosophila* ovaries.

We treated adult flies with 500 μM cytochalasin D, a potent inhibitor of actin polymerization, and analyzed the resulting effects on cytoophidium dynamics and assembly. Live-cell imaging and immunofluorescence analysis revealed that actin filament disruption significantly impaired the dynamic behavior of cytoophidia. Specifically, actin depolymerization arrested the morphological remodeling of cytoophidia, locking them into static, unchanging architectural states (**Figure 6A; Video S6**). This phenotype closely resembled the effects of microtubule disruption, suggesting that both cytoskeletal systems are critical for maintaining the structural plasticity of cytoophidia.

**Figure 6.**
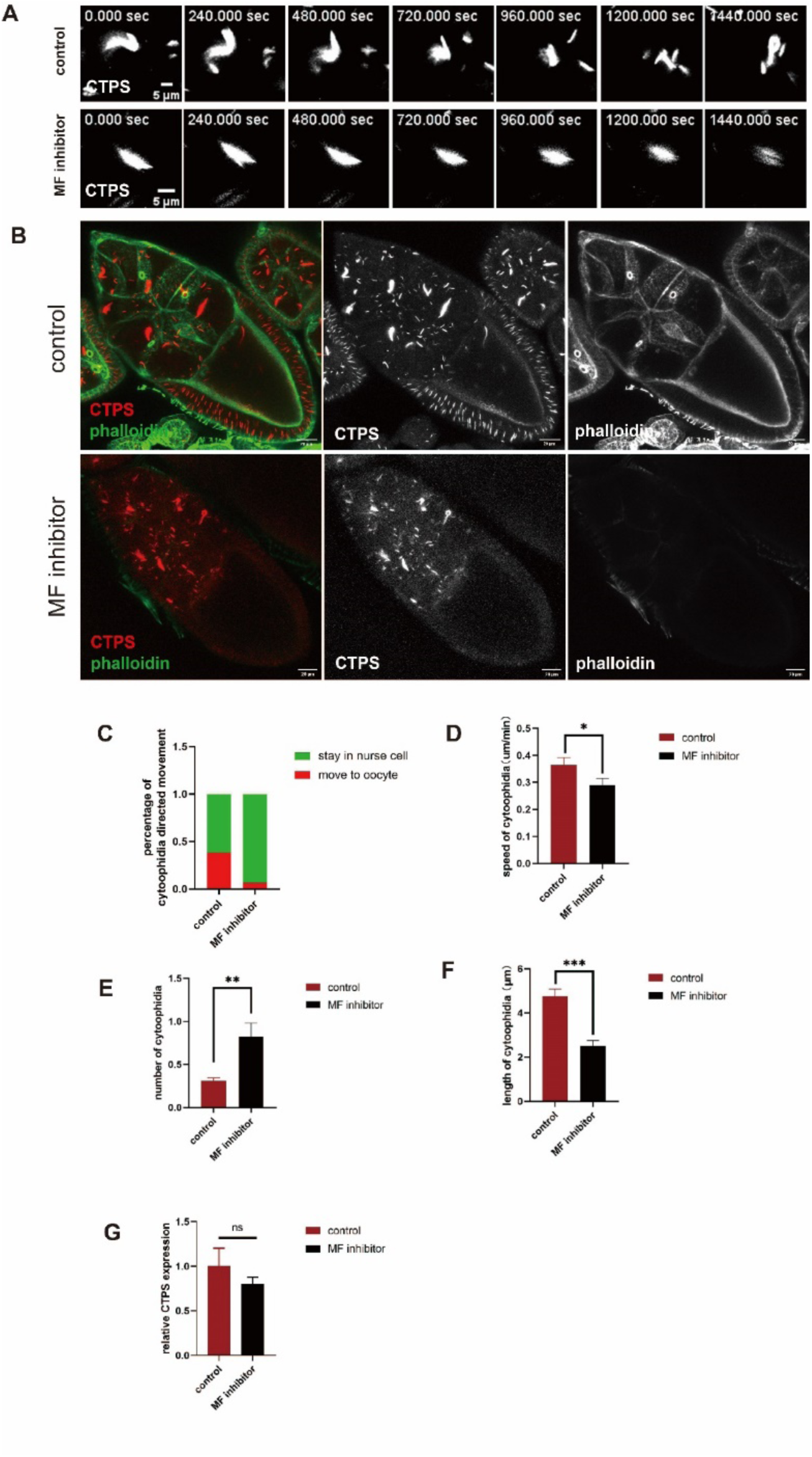
Microfilaments are essential for movement, morphological changes and assembly of cytoophidia. (A) Time-lapse imaging revealed cytoophidia morphological changes, The time marked in the upper left corner represents the time since the capture began. The CTPS (gray)signal shown is obtained using mCherry-tagged CTPS. Scale bar = 5μm. (B) Cytoophidia abundance and length are affected by microfilaments depolymerization. Microtubules (green) are labeled with fluorescein isothiocyanate [FITC]-conjugated tubulin antibody, CTPS (red)signal shown is obtained using mCherry-tagged CTPS. scale bars, 20μm. (C) Quantification of cytoophidia transport through nurse cell-oocyte ring canals in control and microfilaments inhibited groups. (D) The speed of cytoophidia movement. We analyzed 48 filaments in control group and 48 filaments in microfilaments depolymerization group, Data are represented as mean ± SEM, p-value = 0.0443, *. (E) Length of cytoophidia in ovary nurse cell. We analyzed 184 filaments in control group and 184 filaments in microfilaments depolymerization, Data are represented as mean ± SEM, p-value = 0.0001, ***. (F) Number of cytoophidia in ovary nurse cell. We analyzed 9 ovaries in control group and 450 filaments in total, and 9 ovaries in microfilaments depolymerization and 835 filaments in total, Data are represented as mean ± SEM, p-value = 0.003, **. (G) Relative CTPS mRNA expression levels as measured by quantitative RT-PCR using ovary from the control and microfilaments inhibitor-treated flies (6 ovaries/group, 3 biological replicates). Data are represented as mean ± SEM, ns, no significance.

In addition to impaired morphological dynamics, actin inhibition also led to the fragmentation of large cytoophidial structures into smaller, disassembled units (**Figures 6B, 6E, 6F**). Notably, quantitative analysis revealed a shift in cytoophidium size distribution toward smaller fragments, indicative of a loss of higher-order assembly. Importantly, this structural disassembly occurred without significant changes in CTPS transcriptional expression, as determined by qPCR (**Figure 6F**). These findings suggest that actin filaments are specifically required for maintaining the integrity and higher-order assembly of cytoophidia, independent of CTPS expression levels.

Furthermore, quantitative analysis of cytoophidium motility and transport behavior demonstrated that actin inhibition led to a dramatic reduction in their intracellular motility. The frequency of directed transport events from nurse cells to oocytes through ring canals was significantly decreased (**Figures 6C, 6D; Video S6**). These results indicate that actin filaments play a critical role in mediating the intercellular trafficking of cytoophidia, likely by facilitating their movement along actin-based structures or through interactions with motor proteins such as myosin.

Taken together, these findings demonstrate that actin filaments are essential for both the dynamic remodeling and higher-order assembly of cytoophidia, and are additionally required for their efficient intercellular transport in the *Drosophila* ovary. The phenotypic similarities between actin and microtubule disruption further suggest that cytoophidium dynamics rely on a coordinated interplay between multiple cytoskeletal systems.

### Myosin II acts as a key motor protein in regulating cytoophidium dynamics and assembly

Myosins are a superfamily of actin-based motor proteins that generate mechanical force through ATP-dependent interaction with actin filaments, enabling the transport of cellular cargoes and the regulation of cytoskeletal dynamics. Among them, myosin II is a key regulator of cellular contractility, motility, and the maintenance of cellular architecture, and is implicated in processes requiring dynamic remodeling of the actin cytoskeleton. To investigate the potential role of myosin II in modulating cytoophidium dynamics, we treated *Drosophila* samples with 500 µM para-nitroblebbistatin, a selective inhibitor of myosin II ATPase activity [40], for 24 hours, and analyzed the effects on cytoophidium behavior.

Our results demonstrated that inhibition of myosin II significantly suppressed the morphological dynamics of a subset of cytoophidia, leading to a noticeable reduction in their ability to undergo shape changes such as bending, twisting, or remodeling (**Figure 7A**). This observation indicates that myosin II contributes to the structural plasticity and dynamic remodeling of cytoophidia, likely through its role in actin-mediated cytoskeletal organization.

**Figure 7.**
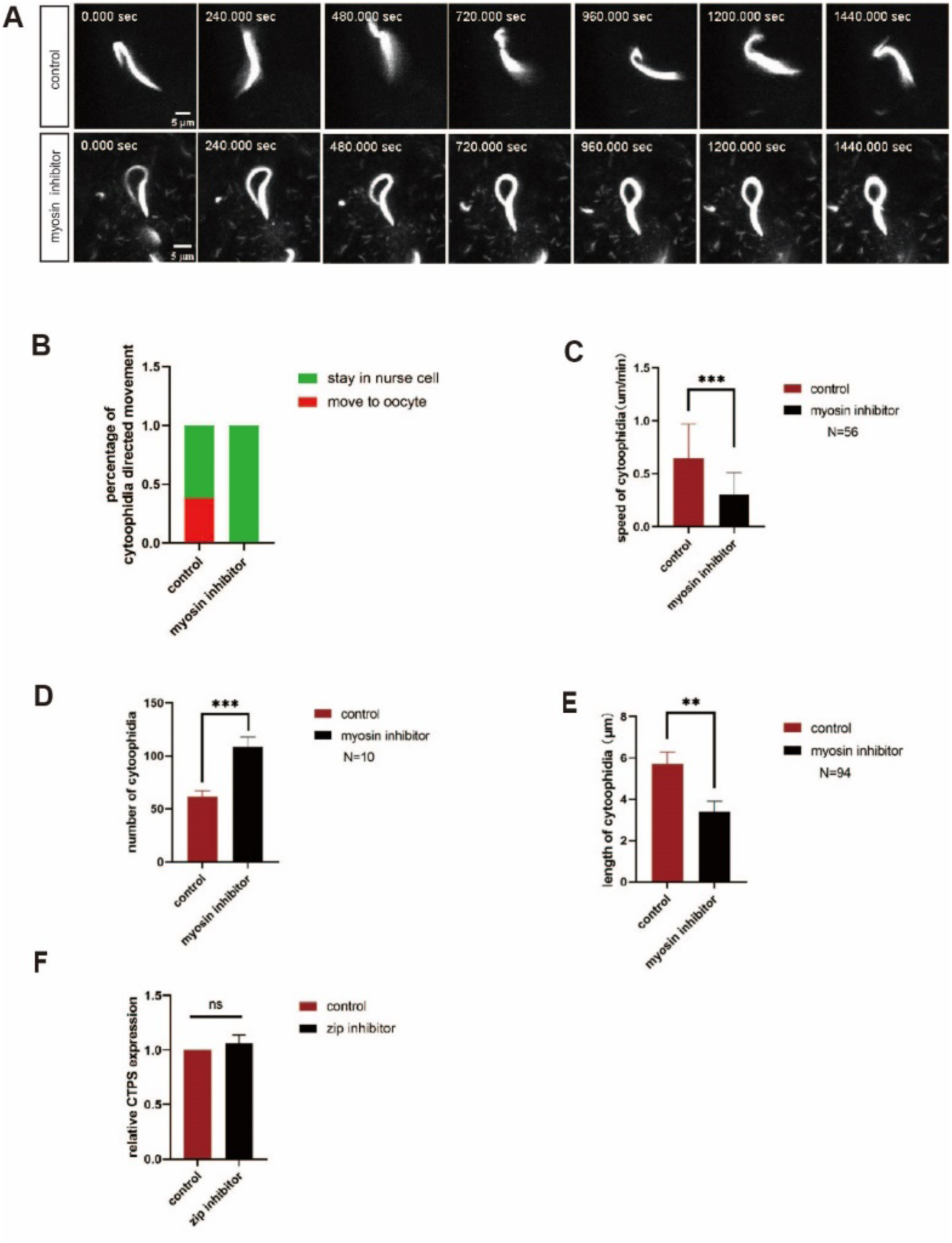
Myosin is essential for movement, morphological changes and assembly of cytoophidia. (A) Time-lapse imaging revealed cytoophidia morphological changes, The time stamps represent the time since the capturing began. The CTPS (gray) signal shown is obtained using mCherry-tagged CTPS. Scale bar = 5μm. (B) Quantification of cytoophidia transport through nurse cell-oocyte ring canals in control and myosin inhibited groups. (C) The speed of cytoophidia movement. We analyzed 56 filaments in control group and 56 filaments in myosin inhibited group, Data are represented as mean ± SEM, ***, p-value = 0.0001. (D) Number of cytoophidia in ovary nurse cell. We analyzed 10 ovaries in control group and 581 filaments in total, and 10 ovaries in myosin inhibited and 989 filaments in total, Data are represented as mean ± SEM, ***, p-value = 0.0005. (E) Length of cytoophidia in ovary nurse cell. We analyzed 94 filaments in control group and 94 filaments in myosin inhibited, Data are represented as mean ± SEM, **, p-value = 0.0040. (F) Relative CTPS mRNA expression levels as measured by quantitative RT-PCR using ovary from the control and myosin inhibitor-treated flies (6 ovaries/group, 3 biological replicates). Data are represented as mean ± SEM. ns, no significance.

In addition to impaired morphological flexibility, myosin II inhibition also markedly reduced the overall motility of cytoophidia. Most notably, we observed a complete loss of intercellular transport of cytoophidia from nurse cells to the oocyte via ring canals (**Figures 7B, 7C; Video S7**). This finding highlights the essential role of myosin II in mediating the directed, intercellular movement of cytoophidia, a process critical for oocyte development.

Furthermore, quantitative analysis of cytoophidium size distribution revealed a significant increase in the abundance of small cytoophidial structures following myosin II inhibition. These smaller cytoophidia were statistically shorter in length compared to those observed in control samples (**Figures 7D, 7E, 7F**). Given that no major changes in CTPS expression were noted in earlier experiments (see Results above and related controls), this phenotype suggests that myosin II is critically involved in the assembly or maintenance of higher-order cytoophidium structures, rather than in the regulation of CTPS levels per se.

Taken together, these findings demonstrate that myosin II plays a multifaceted role in the regulation of cytoophidium biology: it is essential for their dynamic morphological remodeling, their intercellular transport, and their proper assembly into functional filamentous structures. These results further emphasize the coordinated action of multiple cytoskeletal motors—including myosin II, dynein, and potentially others—in orchestrating the complex behavior of cytoophidia within the *Drosophila* ovary.

## DISCUSSION

### Cytoskeletal control of cytoophidium assembly and morphological plasticity

Our findings reveal that the cytoskeleton plays an essential and multifaceted role in regulating both the assembly and dynamic morphology of cytoophidia. Pharmacological disruption of microtubules using colchicine led to a marked reduction in large, filamentous cytoophidia and a concomitant increase in smaller cytoophidial fragments, suggesting that microtubules are required to maintain the higher-order assembly and structural integrity of these membraneless organelles. Similarly, inhibition of actin filament polymerization with cytochalasin D resulted in a comparable phenotype, characterized by impaired structural remodeling and the emergence of smaller cytoophidial units. These observations indicate that both microtubules and microfilaments are critical for sustaining the structural stability and dynamic plasticity of cytoophidia, although their precise contributions may differ in mechanism or context.

These findings resonate with previous biochemical and structural studies of CTPS, the enzymatic core of cytoophidia, which polymerizes into filamentous assemblies both in vivo and in vitro. CTPS forms tetramers that serve as its basic functional units CTPS [41–43], and in Drosophila, these tetramers further assemble into filaments through interactions within the α-helix region of the GAT domain [44]. In parallel, recombinant Escherichia coli CTPS (ecCTPS) has been shown to form bundled filamentous structures under controlled conditions in vitro [45], supporting the idea that CTPS has an intrinsic propensity to self-assemble into higher-order structures. However, the mechanisms by which these CTPS filaments or bundles mature into the large, light-microscopy-visible cytoophidia observed in vivo remain poorly understood.

Our data strongly support the hypothesis that the cytoskeleton—particularly microtubules and actin filaments—mediates the transition from CTPS polymerization to functional cytoophidium assembly. This may occur through mechanisms such as spatial scaffolding, confinement, or active transport that facilitates higher-order aggregation. Nonetheless, the precise molecular determinants of cytoophidium assembly, including the signals and adaptors that link CTPS filaments to the cytoskeleton, as well as the rules governing their size, distribution, and coalescence, remain to be elucidated.

### Cytoskeletal motors orchestrate cytoophidium dynamics and directed transport

A central discovery of our study is the essential and specialized role of cytoplasmic dynein in mediating the long-range, directional transport of cytoophidia from nurse cells to oocytes. Inhibition of dynein with ciliobrevin D abolished cytoophidium trafficking through ring canals, while leaving their intrinsic motility, remodeling capacity, and speed largely unaffected. This functional dissociation underscores that dynein is not broadly required for cytoophidium dynamics per se, but is specifically dedicated to their directional, intercellular transport—a process critical for oocyte provisioning.

In vivo, the activity of dynein depends on its assembly into a multi-protein transport complex that includes dynactin and specific adaptor proteins, which together ensure precise motor-cargo recognition and processive transport along microtubules [46–48]. In *Drosophila* oogenesis, a highly conserved dynein-dynactin-based machinery, also involving Bicaudal D (BicD) and Egalitarian (Egl), drives the polarized transport of mRNP complexes, organelles, and proteins from nurse cells to the oocyte [49–52]. This system ensures that the transcriptionally silent oocyte receives essential developmental determinants via microtubule-guided transport.

For cytoophidia—as non-membrane-bound, metabolically active organelles—the following key questions emerge:

1. How are cytoophidia selectively recognized and recruited by dynein (and potentially other motors) in a spatiotemporally regulated manner?
2. What molecular or cellular cues trigger their dynamic remodeling, assembly, or transport?

Addressing these questions will be essential for elucidating the mechanistic basis of organelle-specific transport and the integration of cytoophidia within broader cellular logistics networks.

### Functional significance of cytoophidium transport and dynamics in oogenesis

Beyond their structural role, the transport and dynamic behavior of cytoophidia likely serve important physiological functions in oogenesis. We propose that cytoophidium dynamics contribute to both metabolic enzyme redistribution and intracellular homeostasis. By shuttling CTPS-containing cytoophidia between nurse cells and the oocyte, the system may facilitate rapid, localized metabolic support, effectively acting as a buffer to stabilize free CTPS concentrations and modulate metabolic flux in response to developmental or environmental cues.

This model is consistent with previous studies showing that CTPS filamentation is responsive to cellular stress and linked to modulation of enzyme activity modulation [45]. Moreover, given that *Drosophila* oocytes are transcriptionally silent during growth and depend entirely on nurse cell-derived cargoes— including mRNAs, proteins, and organelles—for their development [53], the transport of cytoophidia via microtubule-based motors is likely to be tightly integrated with oocyte metabolic regulation and developmental progression.

Thus, cytoophidium dynamics are not merely a reflection of passive CTPS polymer behavior, but are functionally coupled to the oocyte’s need for metabolic coordination, resource allocation, and intercellular communication. Future studies aimed at dissecting the molecular signals governing their transport will shed light on how these organelles contribute to oocyte maturation and reproductive success.

## Conclusion

In summary, our study demonstrates that the cytoskeleton—through the coordinated action of microtubules and microfilaments—plays a central role in regulating the assembly, structural dynamics, and directional transport of cytoophidia in the *Drosophila* ovary (**Figure 8**). We identified dynein as the primary motor protein responsible for long-range, intercellular cytoophidium trafficking, while myosin II was shown to regulate their morphological plasticity and local assembly. Kinesin may also contribute to their transport, though its role appears more limited or redundant under physiological conditions.

**Figure 8.**
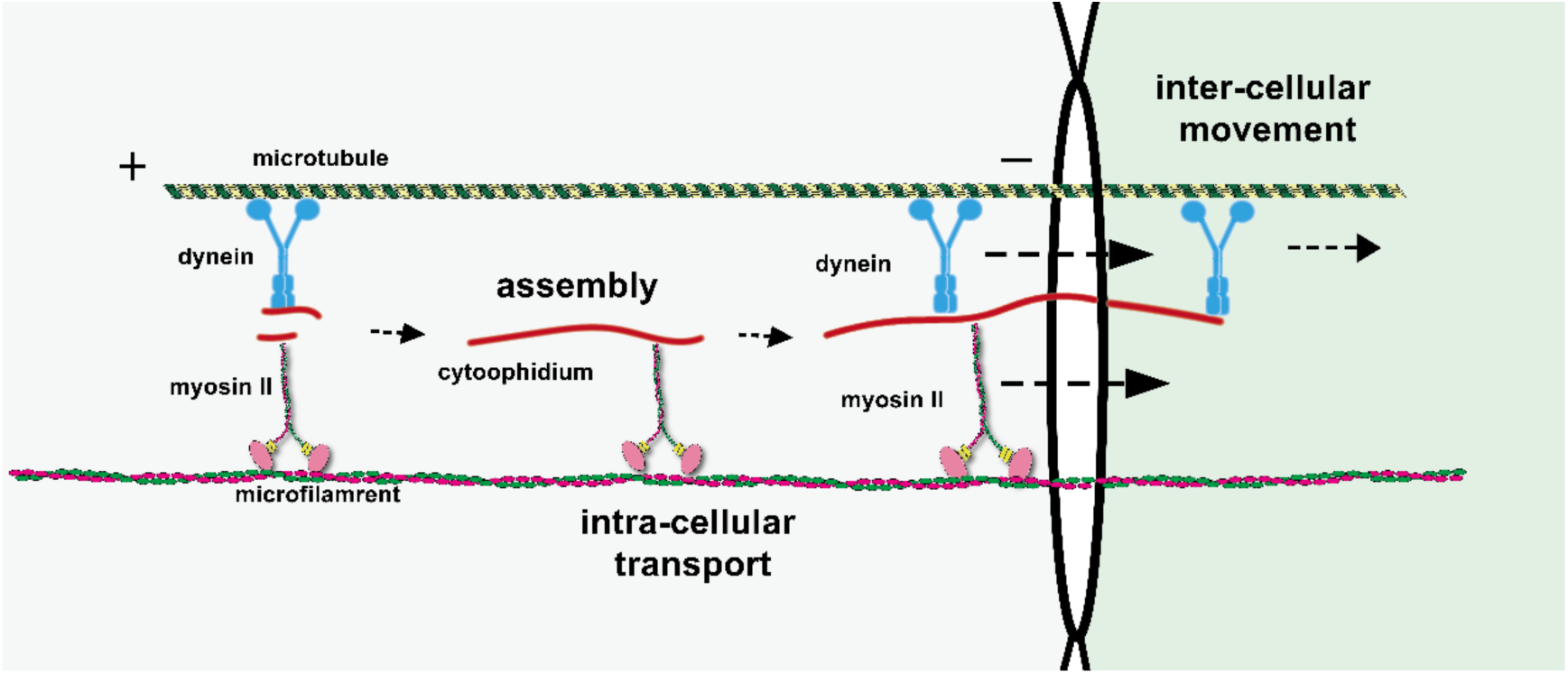
Dynamic Regulation Model of Cytoophidia Dynamics. The diagram of a working model for cytoophidia dynamics. Under physiological conditions, the morphology of cytoophidia is variable. In addition, cytoophidia exhibit two distinct modes of transport: intracellular movement and intercellular movement. Microfilament and microtubules mediate the active transport of cytoophidia. Dynein and myosin are responsible for inter-cellular transport of cytoophidia from the nurse cells to the oocyte.

These findings highlight the remarkable complexity and specificity of cytoskeletal regulation in controlling the lifecycle of a membraneless organelle. They also provide a foundation for future investigations into the molecular mechanisms underlying cytoophidium recruitment, assembly, and transport, as well as their broader physiological roles in oogenesis and beyond.

## ACKNOWLEDGMENTS

We thank the Molecular Imaging Core Facility (MICF) and Molecular Cell Core Facility (MCCF) at the School of Life Science and Technology, ShanghaiTech University for providing technical support. We thank the Core Facility of Drosophila Resource and Technology, CEMCS (Center for Excellence in Molecular Cell Science), CAS (China Academy of Science).

## STATEMENT AND DECLARATIONS

## CONFLICTS OF INTEREST

The authors declare no conflicts of interest.

## AUTHOR CONTRIBUTIONS

X.-J. Liu: Conceptualization, Methodology, Investigation, Formal Analysis, Writing—Original Draft, and Visualization. Y.-L. Li: Conceptualization, Advising, Writing—Review and Editing. S.-Y. Pang: Investigation. K. Dou.: Resources, Writing—Review, Advising. J.-L. Liu: Conceptualization, Resources, Writing— Review and Editing, Supervision, and Project Administration. All authors have read and agreed to the published version of the manuscript.

## FUNDING

This work was supported by the grants from the Ministry of Science and Technology of China (No. 2021YFA0804700), National Natural Science Foundation of China (Grant Nos. 32370744 and 32350710195), and UK Medical Research Council (Grant Nos. MC_UU_12021/3 and MC_U137788471) for grants to J. L. L. and National Natural Science Foundation of China (Grant No. 32370604) to K.D.

## DATA AVAILABILITY

All data are available in the text and supplementary information. Original data are available upon request

## Notes

### Competing Interest Statement

The authors have declared no competing interest.

