## Supplemental Video legends for "Cytoskeleton Regulates Cytoophidium Dynamics in *Drosophila* Ovaries"

**Corresponding:**

**Video S1.	Three types of dynamic changes in cytoophidia**, **related to Figure 1.**

**Video S2.	Microtubules are essential for cytoophidia morphological changes movement, and assembly., related to Figure 2.**

**Video S3.	Microtubules stablization does not affect motion, morphological changes and assembly**, **related to Figure 3.**

**Video S4.	Dynein is indispensable assembly and directed movement of cytoophidia**, **related to Figure 4.**

**Video S5.  Kinesin does not affect cytoophidia dynamic changes**, **related to Figure 5.**

**Video S6. Microfilaments are essential for movement, morphological changes and assembly of cytoophidia**, **related to Figure 6.**

**Video S7. Myosin is essential for movement, morphological changes and assembly of cytoophidia**, **related to Figure 7.**
